# A Computational Model of Stereoscopic Prey Capture in Praying Mantises”

**DOI:** 10.1101/2021.12.03.471070

**Authors:** James O’Keeffe, Vivek Nityananda, Jenny Read

**Affiliations:** Dyson School of Design Engineering, Imperial College, London, SW7 2AZ, UK; Biosciences Institute, Newcastle University, Newcastle, UK

## Abstract

We present a simple model which can account for the stereoscopic sensitivity of praying mantis predatory strikes. The model consists of a single “disparity sensor”: a binocular neuron sensitive to stereoscopic disparity and thus to distance from the animal. The model is based closely on the known behavioural and neurophysiological properties of mantis stereopsis. The monocular inputs to the neuron reflect temporal change and are insensitive to contrast sign, making the sensor insensitive to interocular correlation. The monocular receptive fields have a excitatory centre and inhibitory surround, making them tuned to size. The disparity sensor combines inputs from the two eyes linearly, applies a threshold and then an exponent output nonlinearity. The activity of the sensor represents the model mantis’s instantaneous probability of striking. We integrate this over the stimulus duration to obtain the expected number of strikes in response to moving targets with different stereoscopic distance, size and vertical disparity. We optimised the parameters of the model so as to bring its predictions into agreement with our empirical data on mean strike rate as a function of stimulus size and distance. The model proves capable of reproducing the relatively broad tuning to size and narrow tuning to stereoscopic distance seen in mantis striking behaviour. The model also displays realistic responses to vertical disparity. Most surprisingly, although the model has only a single centre-surround receptive field in each eye, it displays qualitatively the same interaction between size and distance as we observed in real mantids: the preferred size increases as prey distance increases beyond the preferred distance. We show that this occurs because of a stereoscopic “false match” between the leading edge of the stimulus in one eye and its trailing edge in the other; further work will be required to find whether such false matches occur in real mantises. This is the first image-computable model of insect stereopsis, and reproduces key features of both neurophysiology and striking behaviour.

## 1 Introduction

Depth estimation is a critical task for many natural organisms and has broad applicability for autonomous systems. Stereopsis, the computation of distance via triangulation between two eyes, stands out as a particularly robust solution for depth estimation. It has evolved independently at least five times: in mammals, birds, amphibians, cephalopods and insects [1, 2].

Most machine stereopsis algorithms tend to draw inspiration from humans: comparing patterns of contrast in the two eyes’ images to calculate a detailed map of disparity across the visual field [3, 4]. This is a computationally expensive process, which in humans appears to involve multiple areas of visual cortex [5], and so the performance of such algorithms is restricted by the availability of computational resources [6]. One way of addressing this limitation would be to draw inspiration from stereo algorithms in animals with smaller brains relative to primates. Insects are particularly interesting animals in which to investigate this given that their brains are orders of magnitude smaller than human brains. Despite this they are capable of complex visual tasks. Praying mantises are the only insects so far proven to use stereopsis for depth estimation [7–9]. Experiments have demonstrated this by modifying the disparity between mantis eyes with prisms [7] and more recently, an “insect 3D cinema” using coloured filters to display separate images to each eye (Figure 1).

**Fig 1.**
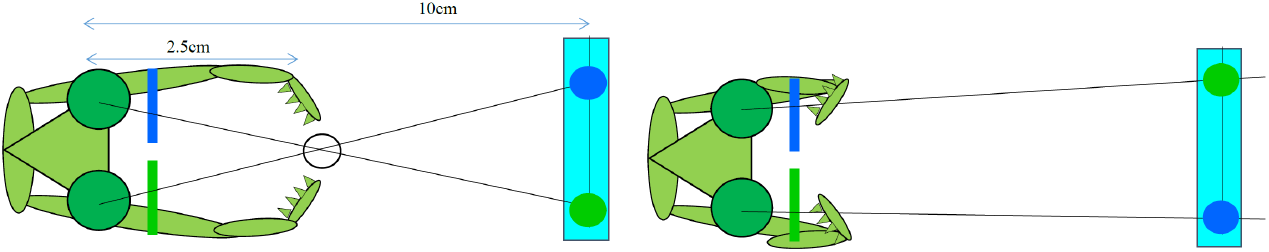
Praying mantis viewing a simulated target in (A) crossed and (B) uncrossed geometry. Each element of the target is displayed on a screen 10 cm away, well outside the catch range. Coloured filters placed over the mantid’s eyes ensure that each eye sees only the intended target. In (A), the target is simulated as being at 2.5 cm, where the lines of sight cross, eliciting a strike. In (B), left and right images are exchanged so the lines of sight diverge. Mantids rarely strike at such stimuli [8, 9].

One way in which mantis stereopsis seems simpler than humans’ is that it probably aims to compute only whether or not a prey item is within catch range, rather than to compute a map of disparity across the visual scene [10]. Additionally, our behavioural experiments have revealed that mantids compute stereoscopic distance using stimulus features different to those used by human stereopsis [11]. Whereas human stereopsis relies on comparing detailed patterns of contrast in each eye, mantis stereopsis looks only for regions of the image where contrast is changing, and is not sensitive to the detailed pattern of contrast. For example, mantis stereopsis is quite happy to accept as “matches” between left and right eyes regions where the contrast is opposite in sign or where local elementary motion is opposite in direction [11, 12].

Intriguingly, however, neurophysiological experiments suggest that the basic computation performed by disparity-selective neurons in the mantis brain may be quite similar to our own [2, 13, 14]. A slightly modified version of the disparity energy model, originally introduced to explain the response of neurons in primary visual cortex of the cat [15] and later applied to primates [16–18], gave a good account of most neurons recorded in the mantis brain. This suggests that mammals and mantids may have independently involved the same basic binocular computation, closely related to cross-correlation [19, 20], although the inputs to this computation may be different: contrast in vertebrates vs temporal change in mantids.

Overall, then, it seems that mantis stereopsis has evolved some of the same techniques as human stereopsis, but uses them within a simpler and computationally cheaper system – yet achieves results that have made the praying mantis a successful ambush predator for millions of years. Given this, a successful model of mantis stereopsis could be beneficial to the field of robotics and artificial intelligence, especially in areas such as swarm robotics, where individual robots are necessarily cheap and lightweight [21]. In this paper we therefore develop a basic model of mantis stereopsis.

Although mantis brains contain multiple distinct classes of disparity-tuned neurons [13, 14], in this first model we include just a single disparity-tuned model neuron or “disparity sensor”, tuned to a single location in space. In reality, even if only a single neuron class were involved in striking behaviour, strikes might reflect activity in several different neurons of this class, e.g. tuned to different locations in space; neuromimetic models of human stereopsis, e.g. [19], invariably assume a population of neurons tuned different disparities. However, it turns out that even a single disparity sensor gives quite a good account of mantis behaviour. Establishing the strengths and limitations of a single-neuron model is an essential prerequisite for understanding the additional contributions of other classes of neuron, as well as for discovering the simplest model that can achieve stereopsis. We therefore examined how well this simple model can capture key aspects of mantis predatory behaviour, including its tuning to prey size and vertical disparity as well as stereoscopic distance or horizontal disparity. We had two specific questions about size and disparity tuning which we expected would challenge a single-sensor model.

The first question is whether a single sensor can account for both the size and the disparity tuning we see in mantids. As well as being sensitive to stereoscopically-defined distance, mantids are also most likely to attack prey of a particular angular size, typically 10-30° [22–26].

In principle, the dependence of mantis striking behaviour on size and disparity could reflect the activity of distinct neurons: some neuron(s) tuned to size but not disparity, and others tuned to distance but not size, both influencing strike probability. However, neurophysiology has revealed the presence of binocular, disparity-tuned neurons with centre/surround receptive fields in each eye [13, 14]. This suggests that such neurons may suffice to explain the tuning of strikes to both size and disparity. The centre/surround receptive field structure could explain the size tuning, with the preferred size corresponding to the central excitatory region of the receptive field. Smaller prey would not fill this excitatory region, and would therefore not maximally activate the neuron, while larger prey would also activate the inhibitory surround, reducing the response. The offset in position of the receptive fields between the two eyes could explain the disparity tuning [15].

However, while it seems clear that such a neuron would be tuned both to size and disparity, qualitatively in agreement with mantis behaviour, it is less clear that it can account quantitatively for the data. Rossel [26] found that strike rate peaked for targets around 25° in diameter, falling to 50% maximal for targets as small as 15 ° or as large as 45°. Thus, the full-width half-maximum bandwidth of size tuning was around 30°. In a slightly smaller species, we found strike rates 50% of maximum for targets from 7 to 25°, i.e. a wide size-tuning bandwidth of around 18°. In contrast, both groups found much narrower disparity tuning. Rossel found that the peak strike rate dropped from 1 for 35mm to 0.6 for stimuli at 55mm, a change of 5° in binocular disparity given the interocular distance of 8mm. We [8] found similar sensitivity, with the strike rate dropping from 80% at 25mm to 20% at 38mm: also a change of around 5° in disparity, given the 7mm interocular distance of our mantids. Thus, the full width half-maximum bandwidth of distance tuning corresponds to around 10°, or an average shift of 5° in each eye.

We therefore wanted to find out whether it is possible to build a sensory neuron whose response stays high over a 20° range of target size, but tolerates only a 5° shift in monocular position, as indicated by the data in Figure 2. This is potentially challenging to achieve, since a neuron tuned to large sizes would necessarily have a large excitatory region and thus be tolerant of small monocular shifts which kept the target largely within this region. If indeed it was not possible to achieve this with a single sensor, this would suggest that mantis striking behaviour might be driven by a combination of several disparity sensors with small receptive fields. The behavioural disparity sensitivity would reflect the receptive field size of individual disparity sensors, while the size preference would reflect pooling of several such sensors with an excitatory-centre/inhibitory-surround pattern of weights. This would be important to know when attempting to unravel the underlying neural circuitry.

**Fig 2.**
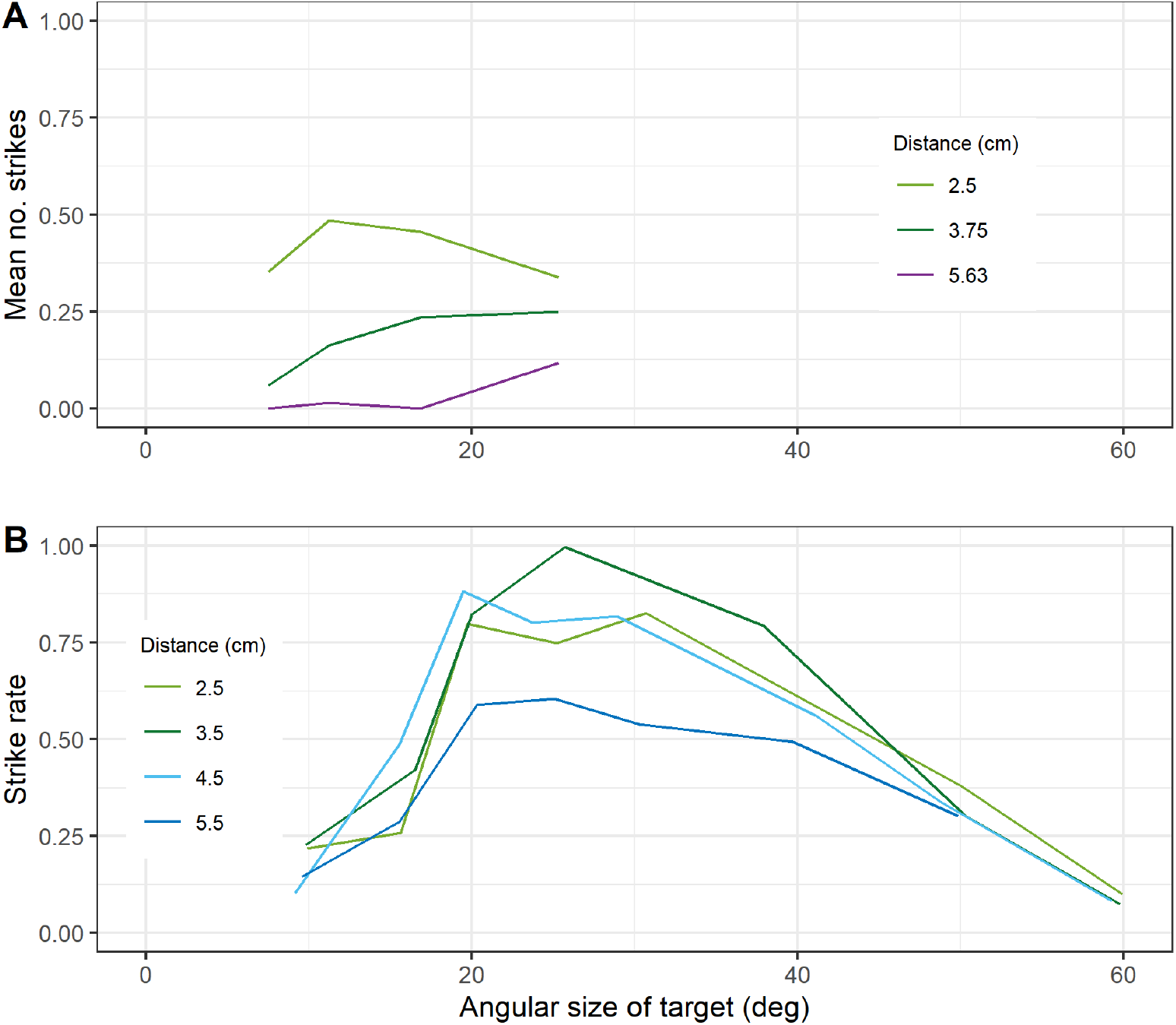
Sensitivity of mantis striking behaviour to stimulus size and stereoscopic distance. (A) Data from [8], available at http://dx.doi.org/10.1098/rstb.2015.0262. Species was Sphodromantis lineola; stimuli were on a computer screen 10cm from the insect; nearer distances were simulated using coloured filters. (B) Data extracted from [26] using WebPlotDigitiser (https://apps.automeris.io/wpd/). Species was Sphodromantis viridis; stimuli were on a computer screen 5.5cm from the insect; nearer distances were simulated using base-out prisms. (A) plots the mean number of strikes per trial; (B) plots the strike rate, i.e. the probability that a given trial elicits striking behaviour, which is slightly different given that some trials elicit two or three strikes. [8]

A second question of interest concerns the relationship between distance and preferred angular size. Size constancy, i.e. a preference for prey of a constant physical size (in cm) regardless of distance, would require a decrease in preferred angular size (in degrees) as the distance increases. Previous work has found no evidence for size constancy [8, 26]. However, while Rossel [26] found that mantids preferred prey of around 25° regardless of distance, we [8] found that preferred angular size actually increased with stereoscopic distance (Figure 2) - the opposite of what would be required for size constancy. It was unclear from previous results why this pattern emerged and the mechanisms underlying it are still unknown. Thus, a second aim of this paper was to understand whether this effect implies the operation of distinct disparity sensors, e.g. one tuned to a stereoscopic distance of 25mm and size 11°, and a second tuned to a distance of 40mm and size 25°. If so, this could provide one possible function for the different classes of disparity-tuned mantis neuron identified by Rosner [13, 14].

To address this, we constructed a model based on a single neuron tuned to both size and disparity, and sought model parameters that could account for our behavioural data.

## 2 Methods

We first describe our model at a conceptual level, before detailing how the simulation was implemented in Matlab.

### 2.1 Model structure

Following our previous work [12], we assume that the images presented to the mantid’s eye first undergo lowpass spatial filtering, representing the response of retinal ommatidia to the pattern of light falling on the eye. They then undergo highpass temporal filtering, representing the response of lamina monopolar cells, which respond mainly to light increments and decrements [27, 28]. Following [29], we used a first-order Butterworth filter with a time-constant of 20ms. We then square the outputs, effectively combining responses to increments and decrements while losing the sign. We denote the result by *J*(*x, y, t*), a function of location on the retina and of time.

In [13], we found that the response of disparity-tuned neurons in the mantis brain was well described with a model based on the disparity energy model originally proposed by [15]. In this model, images are convolved with linear receptive fields in each eye. The output from each receptive field is

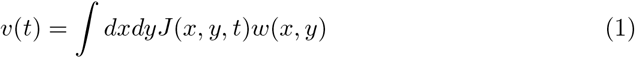

where *J*(*x, y, t*) is the filtered input from that eye, as a function of location and time, and *w*(*x, y*) is the receptive field function – effectively, the synaptic weight connecting input from location (*x, y*) to the disparity-tuned neuron. The instantaneous response of the disparity-tuned neuron is then modelled as

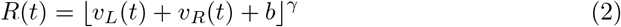

In our fitted model, the bias *b* is negative and so acts as a threshold: the input *v_L_* + *v_R_* must exceed b in order for the response *R* to be non-zero. A positive value of b would model a tonic response, i.e. the cell would be active spontaneously even in the absence of input.

We assume that the response of this neuron represents the instantaneous probability that the mantis will strike. Thus, the mean number of strikes released to a particular stimulus is obtained by integrating *R*(*t*) over the duration of the stimulus:

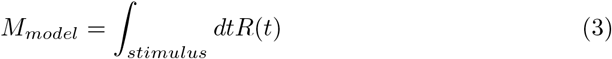

As described below, we adjust the model parameters so as to ensure good agreement between this expected number of strikes, predicted by the model, and the number of strikes observed empirically.

### 2.2 Receptive field structure

The receptive field structure was assumed to be identical in the two eyes but offset horizontally; the amount of this offset controls the disparity tuning of the binocular neuron. We assumed that the receptive field in each eye had an excitatory centre and an inhibitory surround; the size of the excitatory region controls the preferred size of the neuron. For simplicity, our model receptive fields are constructed from square regions of uniform weight (Figure 3). We found that we needed to include both a central excitatory region with high positive weight and an outer excitatory region with lower positive weight, as well as an inhibitory surround with negative weight.

**Fig 3.**
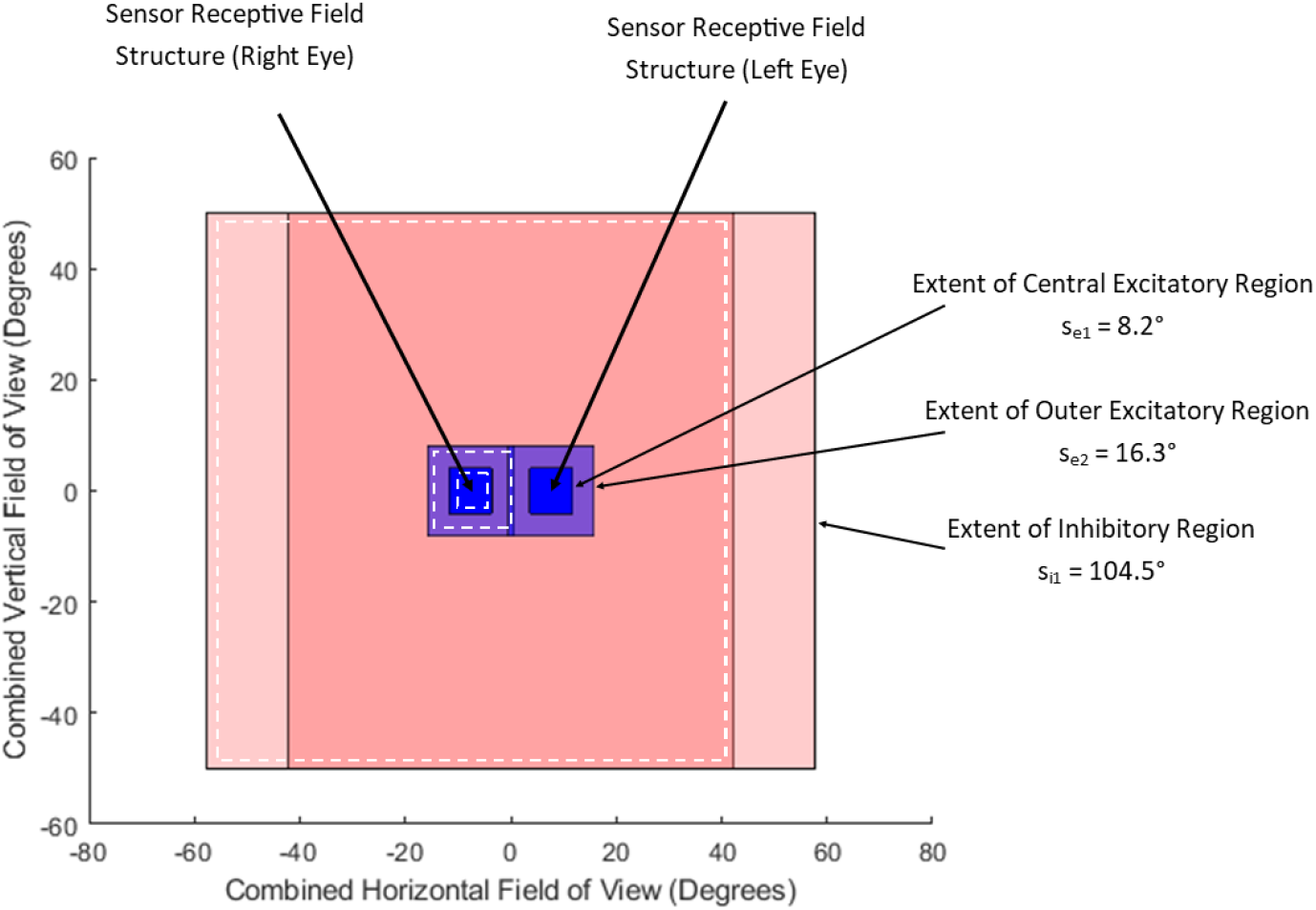
The image shows left- and right-eye receptive fields, superimposed and backprojected onto a screen at 10cm distance for ease of comparison with the stimuli. The left- and right-eye receptive fields have identical structures, with a central strongly excitatory region (blue) surrounded by an outer, more weakly excitatory region (purple), and then by a much larger inhibitory region (pink). The receptive fields in the two eyes are offset horizontally by the screen disparity d = 15.4°.

Considering a monocular target centred on the receptive field (Figure 3), we can see that as the target size increases from zero, the total response will increase rapidly until the target fills the central, strongly excitatory region, and thereafter increase less rapidly until the target fills the outer excitatory region. As the target gets larger still, the response will decrease due to inhibition from the surround. When the target is large enough, inhibition from the surround will cancel out excitation from the centre, and there will be no output (Equation 2).

### 2.3 Parameter optimisation

The model parameters are shown in Table 1. Those listed as “fixed” were set at the specified value, based on previous literature (or pilot experiments in the case of *s_i_*). Those listed as “optimised” were obtained by a maximum likelihood fit, comparing the expected number of strikes to the number empirically observed, assuming Poisson statistics. That is, we sought the set of parameters **p** which maximised the function Equation 4:

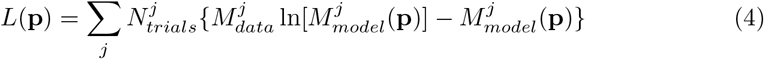

where the sum is over all stimulus conditions (varying in stereoscopic distance and size); 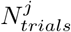 is the number of trials conducted for that stimulus condition; 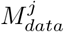 is the mean number of strikes elicited per trial, averaged over all 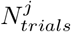 trials; and 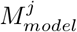 (**p**) is the expected number of strikes per trial predicted by the model (Equation 3) for a stimulus at that distance and size, given the current set of model parameters **p**. This expression is derived from the likelihood of observing *M_data_* given a Poisson distribution with mean *M_model_*, though note that *L* is not exactly equal to the log-likelihood, since for simplicity we have omitted constant terms that do not depend on the model parameters and thus do not affect the optimisation. *N_trials_* would be relevant if it varied between conditions, although in our data-set it was a constant. As expected, *L* is maximised as *M_model_* approaches *M_data_* for every trial.

**Table 1.** Parameters controlling the behaviour of our model, and how these were obtained. The top two rows are for the early visual system, and the remainder describe the disparity sensor. Parameters optimised by fitting to the data were constrained to lie between the stated bounds. The receptive field regions were squares, with the sizes s_e1_, s_e2_, s_i_ being the side-length. i is simply chosen to be very large compared to the stimulus. In our simulation, each pixel represents 0.154 degrees visual angle. The sensor disparity is given both as a screen disparity α_pref_, and in parentheses as the corresponding distance D_pref_.

| Symbol | Meaning | How deter-<br>mined | Lower<br>bound | Upper<br>bound | Value |
| --- | --- | --- | --- | --- | --- |
| N/A | standard deviation of Gaussian lowpass filtering | Fixed |  |  | 0.6° |
| $\tau$ | time-constant of highpass filter | Fixed | | | 20 ms |
| $\alpha_{pref}$ | sensor position screen disparity, i.e. offset left- and right-eye receptive field centres, as measured on a screen at 10cm distance | Optimised | 9°<br>(3cm) | 23°<br>(1.5cm) | 15.4°,<br>(2.1cm) |
| $s_{e1}$ | size of receptive field central excitatory region | Optimised | 3° | 11° | 8.14° |
| $s_{e2}$ | size of receptive field outer excitatory region | Optimised | 11° | 17° | 16.3° |
| $s_i$ | size of receptive field inhibitory region | Fixed | | | 104.5° |
| $w_{e1}$ | weight of receptive field central excitatory region | Optimised | 0 | $+\infty$ | $6.77 \times 10^{-4}$ |
| $w_{e2}$ | weight of receptive field outer excitatory region | Optimised | 0 | $+\infty$ | $3.18 \times 10^{-4}$ |
| $w_i$ | weight of receptive field inhibitory region | Optimised | $-\infty$ | 0 | $7.46 \times 10^{-5}$ |
| $b$ | tonic inhibition onto disparity sensor | Optimised | $-\infty$ | 0 | -0.0542 |
| $\gamma$ | exponent of disparity sensor output power nonlinearity | Optimised | 0 | $+\infty$ | 5.05 |

### 2.4 Stereoscopic distance and definitions of disparity

Our simulations are closely based on our behavioural and neurophysiological experiments, in which stimuli were presented on a screen 10cm from the animal’s eyes, as shown in Figure 4. Accordingly, we represent stimuli in the simulations as if they were being presented on a screen a distance *S*=10cm away. Targets at a stereoscopic distance *D* are simulated by giving their images a horizontal separation *P* on the simulated screen. Since we mainly measure image locations in degrees, we also express *P* via the angle *α* that it subtends at the eyes. We refer to this as *screen disparity*:

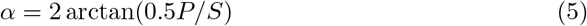

**Fig 4.**
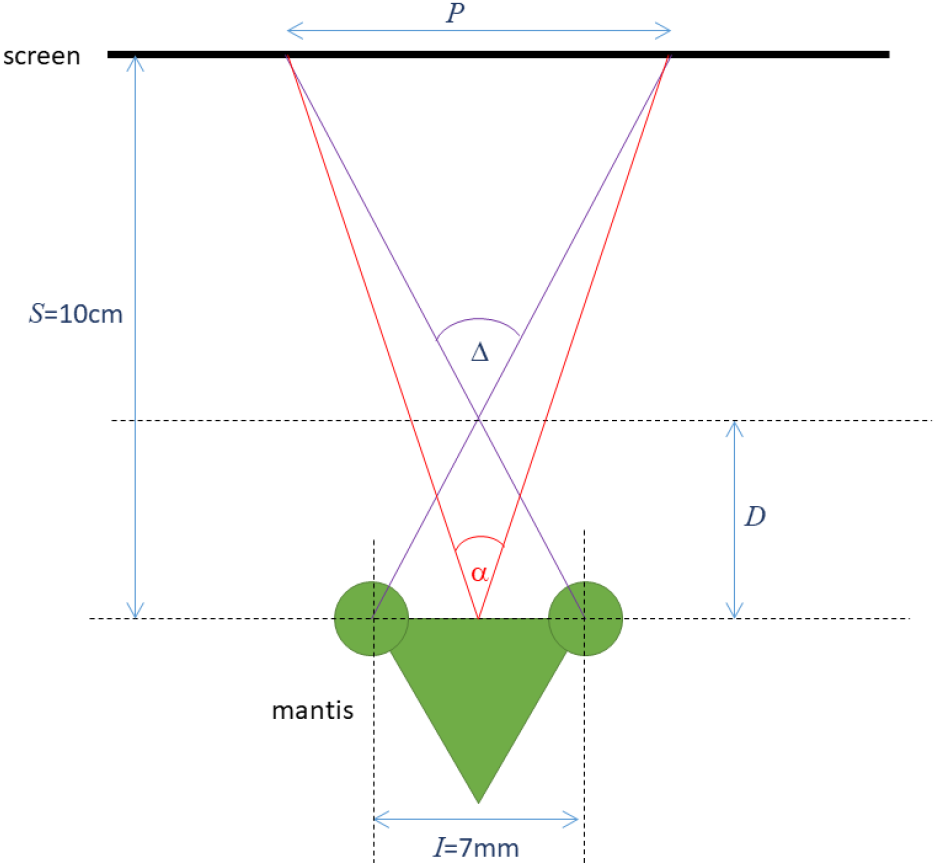
Top-down view (not to scale) of a mantis viewing a computer screen at a distance S. To simulate an object at distance D from the mantis, the left and right eye’s images need to be separated by a distance P on the screen. The angle subtended by the distance P at the mantis is the screen disparity α; the retinal disparity Δ of the virtual object is the angle subtended by the interocular distance I at the virtual object.

In our plots, we will display simulated images as they would have been presented on the screen. The separation between left- and right-eye images will thus correspond to the screen disparity *α*. A screen disparity of *α*, on a screen a distance *S* away from the animal simulates a virtual object at a distance *D*, where

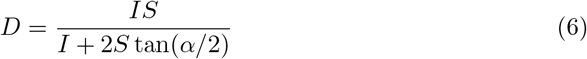

and thus

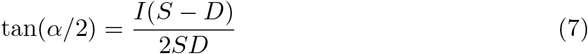

*I* is the interocular separation, which we take to be 7mm throughout this paper. Biologically, the screen disparity *α* is not very relevant, since it reflects where we happened to position our screen in the experiments. A more relevant quality is the angle Δ, the *retinal disparity*. An object at distance *D* from the animal has retinal disparity

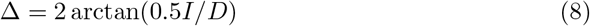

The relationship between screen disparity *α* and retinal disparity Δ is given by

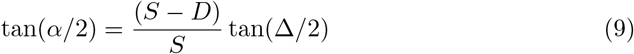

i.e. they become equal when the screen is at infinity. In this paper, we will mainly report the screen disparity *α* and the simulated distance *D*. To recover retinal disparity Δ, use the above expressions with our simulated interocular distance *I*=7mm and screen distance *S* =10cm.

### 2.5 Stimuli

#### 2.5.1 Simulated behavioural experiments

For optimising model parameters and exploring the behavioural response, we use stimuli consisting of bright disks on a dark background (in fact mantids prefer dark stimuli on a bright background, but the model presented here is only sensitive to changes in contrast, not the direction of the change, and so this does not affect our results). In most of our behavioural experiments, including the source of the data used for fitting [8], we used stimuli which moved in a spiral trajectory from the outer edges of a monitor screen to the centre. The target began more than 100° away from the centre of the screen, moving with a velocity of almost 1500° per second, and gradually decelerates over 5 seconds, ending with a velocity of 0 in the centre of the screen.

This stimulus is ill-suited for understanding the properties of our disparity-tuned model neuron, since the response will be affected by the speed and direction of the stimulus, both of which are constantly changing in the spiralling stimulus. In this paper, we therefore chose to examine targets moving with a constant speed either horizontally or vertically. These two cardinal directions are the most important to examine, since the binocular receptive fields are offset horizontally but aligned vertically, and so the response is expected to be different for these cases. For our simulations, we chose a constant stimulus speed of 82° per second. Previous work has shown that this speed is close to optimal for eliciting mantis strikes [22], and we have successfully elicited strikes with stimuli moving horizontally at this speed [12]. The spiralling stimulus of [8] reached this speed when it was 15° from the centre of the monitor, in the region where strikes were most likely to be elicited.

### 2.6 Empirical data

We optimised model parameters using the data from our earlier paper, [8], which is available at http://dx.doi.org/10.1098/rstb.2015.0262. The data for stimuli simulating distances within catching range were shown in Figure 2. In the paper, as a control, we also presented stimuli with equivalent uncrossed disparity, as in Figure 1B. Uncrossed stimuli were identical to stimuli used to depict a given virtual distance, except that left and right images were swapped. In uncrossed stimuli, the strike rate was unsurprisingly very low. However, a strike rate of around 10% was observed for the largest stimuli (25° diameter). Because striking was independent of binocular disparity in uncrossed stimuli, we concluded that this was most likely a defensive behaviour, aimed not at catching prey but at deterring a potential threat, and triggered by the angular size of the stimulus rather than its distance. We would therefore not expect such strikes to be under the neural control of a sensor designed to detect prey within catch range, which is what we are trying to simulate here.

Accordingly, we corrected the raw number of strikes by subtracting strikes in the uncrossed condition from strikes in the equivalent crossed condition. The results are shown in Figure 5.

**Fig 5.**
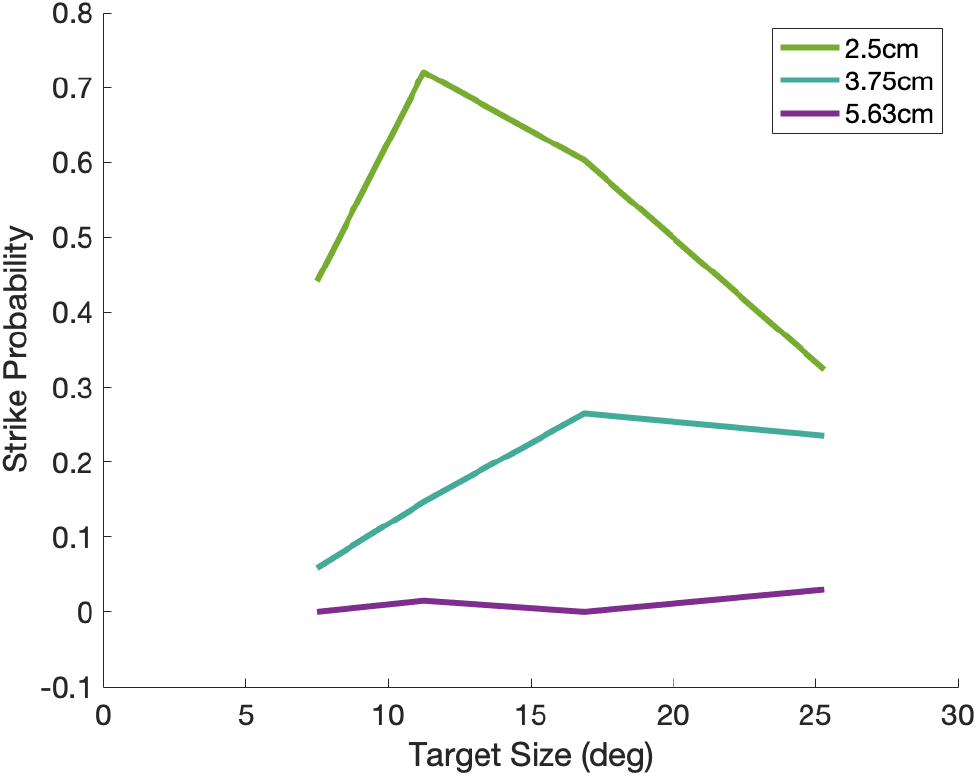
Modified behavioural response of mantises. Mean number of strikes in response to different simulated distances plotted as a function of the angular size of the simulated target in the crossed condition with corresponding strikes in the uncrossed condition subtracted. [8]

For optimising model parameters, we also included synthetic data for monocular, distant and large targets which were not included in our experiments precisely because in our experience they elicit very few strikes (Table 2). These conditions were assigned a strike rate of 0, forcing the optimisation to avoid parameter sets which would predict implausibly large values there. *N_trials_* was set equal to 68 for these conditions too, giving them the same weight as the other conditions.

**Table 2.** Values of M_data_ (mean number of strikes per trial) used for optimisation. Columns show stimulus diameter; rows show stimulus stereoscopic distance. For the top 3×4 cells (distances 2.5-5.63cm and sizes less than 38°), data was derived from [8] by taking the difference between the total number of strikes recorded for crossed vs uncrossed trials for the given stimulus, divided by the total number of times each stimulus was presented to a mantis (N_trials_ = 68: 10 trial presentations on 6 mantids and six presentations on 1 mantis). Zeros for size = 38°, distance = 10cm and monocular stimuli are synthetic data included to force the optimisation to choose parameter values which predict very low strike rates for these conditions, in keeping with our observations. [8]

|  | 7.5° | 11.25° | 16.88° | 25.31° | 38.00° |
| --- | --- | --- | --- | --- | --- |
| 2.50cm | 0.44 | 0.72 | 0.60 | 0.32 | 0.00 |
| 3.75cm | 0.06 | 0.15 | 0.26 | 0.24 | 0.00 |
| 5.63cm | 0.00 | 0.02 | 0.00 | 0.03 | 0.00 |
| 10cm | 0.00 | 0.00 | 0.00 | 0.00 | 0.00 |
| Monocular | 0.00 | 0.00 | 0.00 | 0.00 | 0.00 |

**Table 3.**
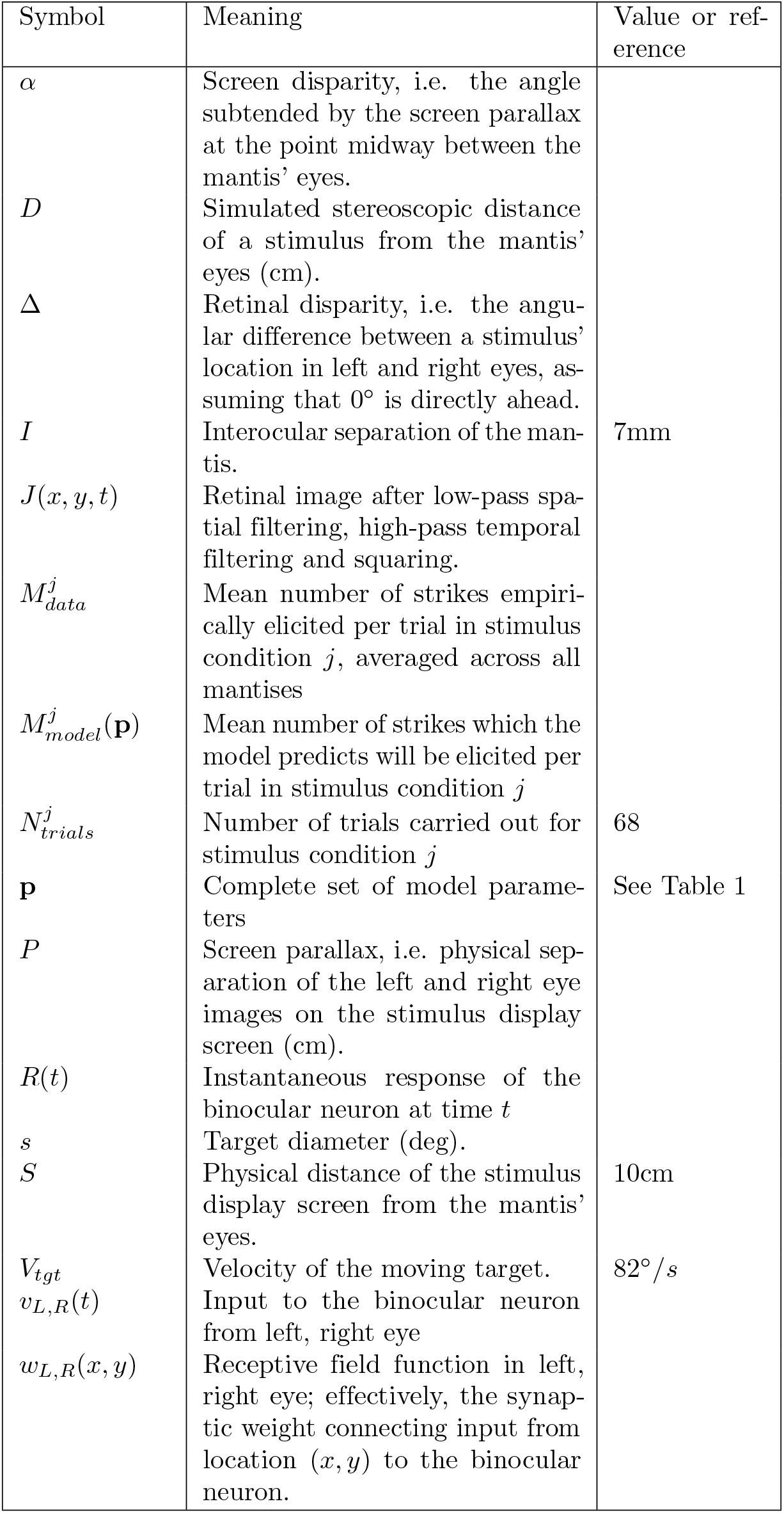
Symbols used in the paper. Symbols used for model parameters are given separately in Table 1, along with values.

As noted above, in our mantis experiments stimuli moved in a spiral trajectory with varying speed and direction, and strikes could be triggered potentially at any point. In our simulations, stimuli moved in a straight line with constant speed. For optimising our model parameters, we used stimuli which passed through the centre of the receptive field (i.e. the average of the left and right receptive fields, given that these were offset horizontally), with zero vertical disparity and a horizontal disparity corresponding to the perceived distance condition. We ran each condition twice: once for vertical motion and once for horizontal. The model’s prediction for the mean number of strikes, *M_model_*, depended on the direction of motion, and we included both conditions in the sum describing our cost function Equation 4, using the same *M_data_* for both as specified in Table 2.

### 2.7 Simulation details

All simulations were carried out in Matlab R2019a (www.themathworks.com). For examining the behavioural responses, we generated the stimuli in each eye as images of 680 × 680 pixels, with each pixel representing approximately 0.15° visual angle. Each stimulus consisted of a light disk (pixel value 1) on a dark background (pixel value 0). The simulation timestep was fixed at 3.3 ms (300Hz), chosen to be substantially lower than the 20ms time-constant of the temporal filter to avoid numerical artefacts. Since the images in our experiments [8] were presented on an LCD monitor with a refresh rate of 60Hz, we updated the images in our simulation every 16.66 ms, i.e. each frame was presented to the model 5 times before the target advanced in the next frame. In each frame update, the target advanced 9 pixels = 1.37° either horizontally or vertically, giving the desired speed of 82°/s.

The horizontal position of the target in each eye was constant for targets moving vertically, and the vertical position was constant for targets moving horizontally. We investigated various choices for these, as discussed in the text. The size of the disk varied between conditions; we used diameters of 3, 6, 9, 12, 16, 19, 22, 25, 28, 31, 34 and 37 degrees. Stimuli crossed the entire visual field of our model, spanning a visual angle of 60°.

These input images were then spatially filtered using the function ‘imgaussfilt’ from the Matlab image processing toolbox, with a Gaussian filter with parameter SD set to 4 (pixels). The SD represents the acceptance angle of the ommatidia, which was approximated at 0.7 deg [30]. These filtered images were then passed through a first-order high-pass temporal filter using Matlab function ‘filter’. Finally, the output of the temporal filter was squared.

These filtered outputs then formed the inputs to the receptive fields of the disparity-tuned neuron. Equation 1 was implemented by point-multiplying each output frame with the model receptive field function in each eye, and summing the result across both dimensions.

After combining the results from both eyes according to Equation 2 at each sampled time-point, Equation 3 was implemented using the Matlab function ‘trapz’. This produced a value of *M_model_* to use in the optimisation cost function, Equation 4. We did this twice for each condition, once for a vertically-moving target and once for horizontally, and added both onto the cost function.

The model parameters in Table 1 were optimised via gradient descent using the Matlab function ‘fmincon’, with –L (Equation 4) as the function to be minimised. fmincon finds the minimum of a constrained nonlinear multivariable function; we constrained all model parameters to have a lower bound of 0 with the exception of the bias parameter *b* (Equation 2), which was constrained to be negative, between -∞ and 0.

The choice of starting point is often critical in multi-dimensional optimisation problems such as this. We used initial values chosen at random within the upper and lower bounds for each parameter (see Table 1). The upper and lower bounds were selected non-randomly, to reflect our own prior knowledge of the sensor where possible. For example, since we know that the strike rate is highest for distances around 2.5cm, we constrained the sensor disparity *d* to correspond to distance in the range 1.5cm to 3cm. In the process of obtaining the parameter values we settled upon and present in this work, the optimisation process was performed many times (before and after the presented parameter set was found) and each parameter with non-infinite upper and lower bounds was tried with starting values spanning the range of the possible values it could assume, to improve the chances of finding the true global optimum.

## 3 Results

### 3.1 Predicted striking behaviour as a function of target size and binocular disparity

Figure 6 shows the predictions of the fitted model (Table 1) for the expected number of strikes per trial, as a function of target size and distance, compared with empirical results (dashed lines). The results are broadly independent of target direction of motion and broadly in line with empirical results despite the simplicity of the model. Although the peak response in the empirical data was at 2.5cm, the optimisation finds the best match with receptive fields whose binocular position disparity corresponds to a distance of 2.1 cm. Accordingly, the curve for 2cm has the highest peak, with curves for nearer and further distances eliciting a lower response, in agreement with mantis behaviour. The asymmetry in the effect of distance on strike probability reflects the geometry: moving from a distance of 2cm to 2.5cm involves moving the target laterally 1.5° in each eye, whereas moving from 2cm to 1.5cm involves a larger shift of 2.6°. This produces a lower response, as more of the target moves into the inhibitory surround of the receptive field. The model also captures the asymmetry in size tuning, with response falling more slowly as target diameter increases beyond the preferred size than when it decreases below it.

**Fig 6.**
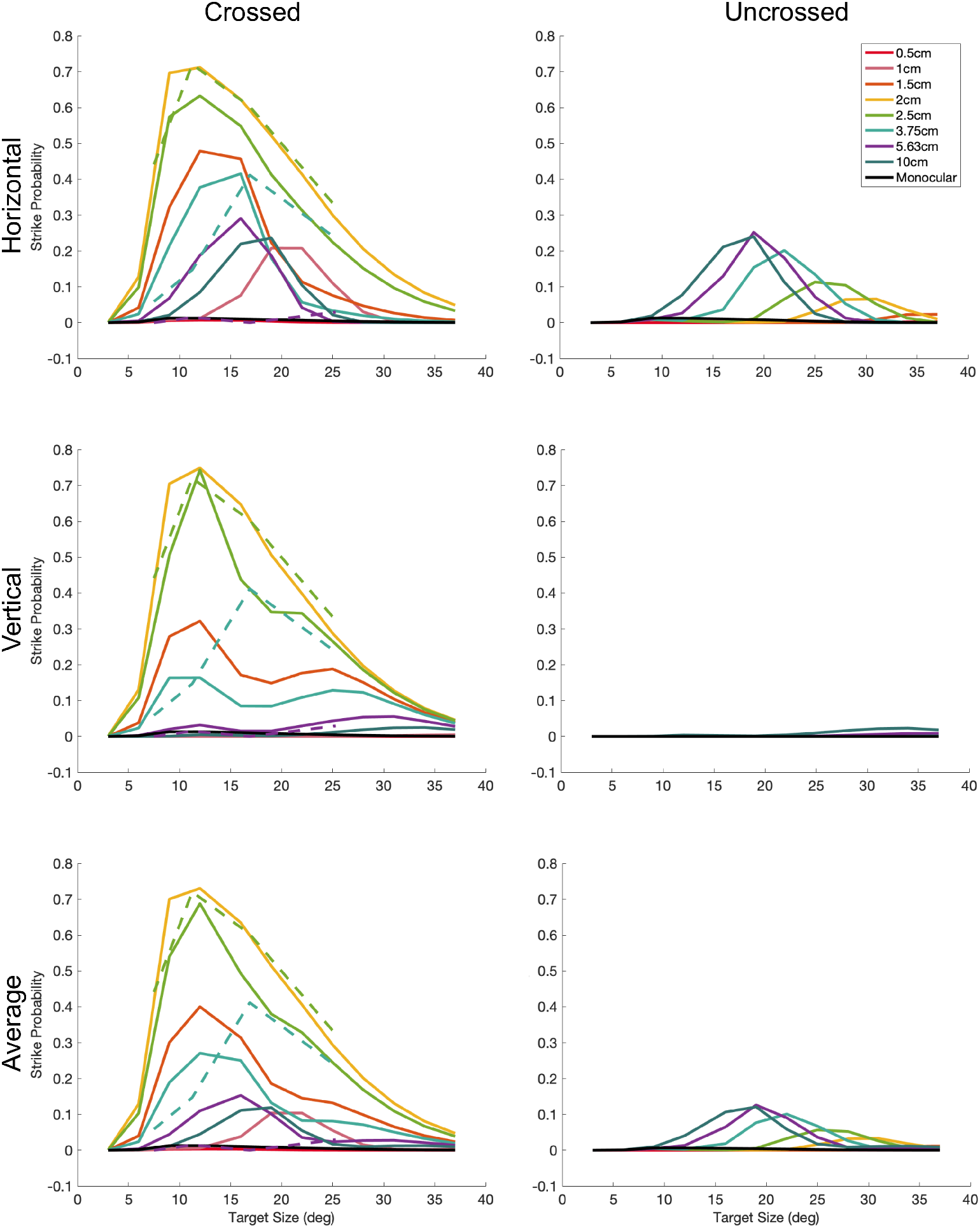
Mean number of strikes per trial predicted by the fitted model, as a function of stimulus size (horizontal axes) and distance (colours). The top row is for targets moving horizontally over the sensor; the middle for stimuli moving vertically, and the bottom row for the average of both. The left panels are for crossed stimuli and the right for uncrossed. In all cases, the target passed directly over the receptive field center. The dashed lines in the crossed panels show the empirical data from [8] (Table 2) which the optimisation aimed to reproduce. In the experiments, the target had a spiralling motion with both horizontal and vertical components, so the same empirical data are shown in all three rows. As described in the Methods (see Table 2), the optimisation also aimed to predict zero strikes for stimuli at 10cm distance, for monocular stimuli, and for stimuli of size 38° diameter. The other conditions - intermediate sizes and distances, and uncrossed stimuli - were not constrained in the fitting, but the model structure ensures that plausible results are also obtained for these.

For targets at the preferred distance of 2.1cm, the predicted strike rate is the same regardless of target direction of motion. However, for other distances, direction of motion makes a difference. This anisotropy reflects the fact that the receptive fields have horizontal, but no vertical, disparity.

Most notably, the model predicts some strikes to large uncrossed stimuli moving horizontally, but virtually none to vertically (cf Figure 6, middle row). No one has yet compared the effect of motion direction on mantis strikes uncrossed stimuli, so we do not know if any such anisotropy occurs empirically. We have previously observed a strike probability of 10-12% for large uncrossed targets ([8], their Fig 5a), but this effect did not depend on target distance, and so, as noted above, we concluded it was not driven by the stereoscopic system.

For horizontal motion, the model also shows an interaction between size and distance tuning. The angular size eliciting the most strikes is 20° for targets 1cm away, then decreases to 12° for targets 1.5cm, then decreases further to 10° for targets 2-2.5cm away, then increases again for more distant targets. The data of [8] showed a qualitatively similar effect.

As we discuss in the next section, the underlying reason for both of these effects is that, for stimuli moving horizontally, it is possible for the leading edge of the disk in one eye to stimulate the sensor at the same time as the trailing edge in the other eye.

### 3.2 Explanation of model behaviour

In any computer model, it is important to understand how the model achieves its results: what features are key to the behaviour (e.g. centre/surround receptive field structure) and which are secondary (e.g. receptive fields modelled as squares). In this section, we dive under the hood of the model in order to understand in detail how it produces its results. Figure 7 shows the response of different components of the model when the virtual stimulus is a disk of size 11° (A), 17° (B) and 26° (C), moving horizontally from left to right across the screen at a simulated distance of 2.5cm. The top row shows the filtered images in left and right eyes, the middle row the inputs to the disparity sensor from left and right receptive fields, while the bottom row shows the response of the disparity sensor.

**Fig 7.**
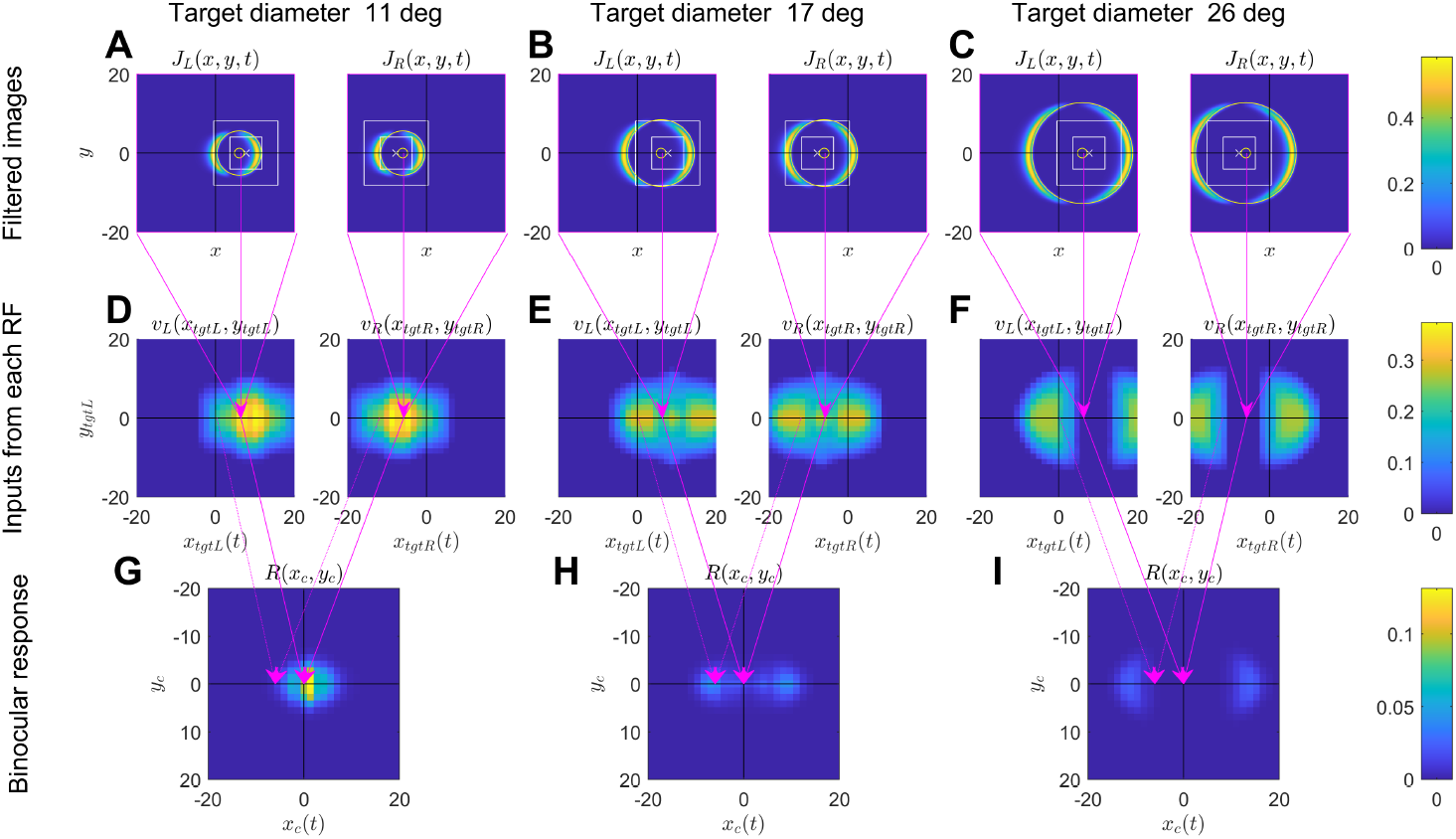
Responses of different model components to a horizontally-moving disk at a simulated distance of 2.5cm from the mantis. Left, middle, right columns are for a target of size 11.2°, 16.9° and 25.5° as indicated. Sub-panels in the top two rows show left and right eyes. In all panels, coordinates are referred to the centre of the screen, i.e. x = 0, y = 0 corresponds to a location 10cm directly in front of the mantis. Top row (ABC): Snapshots of the filtered images J_L,R_(x, y, t), shown as a function of retinal location (x, y) for one particular time t. These are the images reaching the sensor’s receptive fields, following lowpass spatial filtering, highpass temporal filtering and squaring in the early visual system. The receptive field excitatory region is shown superimposed for comparison. Each pixel represents the value of the filtered image at a particular location in the retina. These snapshots are for one particular time t and thus for one particular target position x_tgt_(t), y_tgt_(t) as the target moves across the screen. In this figure, the target was moving horizontally, so y_tgt_ is in fact independent of time whereas x_tgt_ = x_0_ + Vt. The yellow circle marks where the center of the target is in that eye at the time shown. The white cross marks the center of the sensor receptive field in that eye; the inner white square marks the boundary of the central excitatory region, while the outer white square marks the boundary of the outer excitatory region. The surrounding inhibitory region extends beyond the range shown in each panel. Thus, parts of the filtered image falling outside the white squares have an inhibitory effect on the sensor. Middle row (DEF): Inputs to the binocular disparity sensor from the two eyes, v_L,R_. The input from each eye is the inner product of the monocular receptive field with the filtered image at that moment in time. It is here represented as a function of target position x_tgt_(t), y_tgt_. Since the target is moving horizontally across the screen from left to right, x_tgt_ is a function of time, whereas y_tgt_ is constant for a given trajectory. Each pixel-row in DEF therefore represents the time-course of the monocular input, v_L,R_(t), as the target moves from left to right over the screen, at the vertical location ytgt corresponding to the height of the pixel row. The pink arrows mark the value of the monocular input in D for the filtered image shown in A. Bottom row (GHI): response of the disparity sensor, Equation 2. Arrows from D to G show the target locations shown in the top row, A. The target’s direction in the visual field is x_c_ = (x_tgtL_ + x_tgtR_)/2 and y_c_ = (y_tgtL_ + ytgtR)/2. Since the target is moving horizontally, x_c_ is a function of time, but y_c_ is constant for a given trajectory.

#### 3.2.1 The filtered images

Before understanding the response of the disparity sensor itself, we first need to understand what’s happening in each eye. The top row of Figure 7 shows snapshots of the filtered image, *J*(*x, y, t*), at the instant where the disk crosses the midline.

Superimposed on this for reference, a large yellow circle marks the edge of the target at the instant shown; the small yellow o marks its centre. The target disk is at a distance 2.5cm in front of the mantis, corresponding to a screen disparity of 12°. At the moment shown, the disk is centered on *x_tgtL_* = 6° in the left eye and *x_tgtR_* = −6° in the right, so that its overall direction is straight ahead (*x_c_* = (*x_tgtL_* + *x_tgtR_*)/2 = 0).

Because of the high-pass temporal filtering, the filtered image appears as two crescents, lagging behind the leading and trailing edges of the disk as it passes over the photoreceptors from left to right. In general, the time that elapses between the leading and trailing edges is *T* = *s/V*, where *s* is the target diameter and *V* its speed. If *T* is large compared to the timeconstant *τ* of the highpass filter, leading and trailing edges will be clearly separated and will have the same amplitude. The width of each edge will be proportional to speed *V*. As *T* decreases, the amplitude of the trailing edge relative to the leading one will decrease as [1 – exp(–*T/τ*)]^2^. Eventually, for *s* < *V_τ_*ln(2) the edges will merge, without any dip between the leading and trailing edges. In this paper, we use a timeconstant of *τ*=20ms and a target speed of *V* = 82° per second. The edges are therefore smeared over roughly 1.64°, and they do not merge except for diameters below 1°.

The central and outer excitatory regions of the disparity sensor’s receptive fields are also superimposed for comparison in Figure 7ABC (concentric white squares). The center of the receptive field is marked with a white cross; it is at *x_L_* = +7.7° in the left eye and *x_R_* = −7.7° in the right, giving it the sensor disparity of *α_pref_* = 15.4°. The disparity sensor is thus tuned to a location 2.1cm directly in front of the mantis. Recall that surrounding the excitatory regions shown is a much larger inhibitory region. The 3.4° relative disparity between the sensor (disparity *α_pref_* = 15.4°) and the target (disparity 12°) explains why the target is at different locations relative to the receptive fields in the two eyes.

#### 3.2.2 Inputs from each eye’s receptive field

As we saw in Equation 1, the instantaneous input from each eye to the binocular sensor, *v_R_*(*t*) and *v_L_*(*t*), is the result of point-multiplying the receptive field function with the filtered image *J*(*x, y, t*) at that instant. Thus, for each subpanel in Figure 7ABC, there will be a single value of *v_L,R_*(*t*) at the time of that snapshot. Its value will of course depend on where the target is located relative to the receptive fields at the moment of the snapshot.

This is illustrated in Figure 7DEF. Here, each subpanel shows the values of *v_L,R_* as a function of the current target position in that eye. The pink arrows linking each subpanel of Figure 7ABC to a pixel in the corresponding subpanel of Figure 7DEF show where the example images shown in Figure 7ABC are represented in Figure 7DEF. Other pixels in the same row of Figure 7DEF would represent the value of *v_L,R_* at different points in the target’s trajectory. Reading across a single pixel row in

Figure 7DEF, we can see how the values of *v_L,R_* change as the disk crosses the receptive field. This is why the x-axis is labelled *x_tgt_*(*t*), since the x-position of the target is changing with time. A horizontal trajectory with a different y-location, higher or lower on the screen, would be represented by a different pixel-row Figure 7DEF, with the appropriate value of *y_tgt_*.

For the smallest disk shown, Figure 7D, there is a single central peak. This occurs when the target is moving across the excitatory regions of the receptive field. At the moment shown in Figure 7A, in the left eye, activation caused by the leading edge of the disk is in the central excitatory region, where excitation is strongest. The trailing edge is mainly in the outer excitatory region, but some is still in the inhibitory region. In the right eye, both leading and trailing edges are mainly in the outer excitatory region, causing weak excitation but no inhibition. As we see from the corresponding pixel values in Figure 7D, the net effect is similar in both eyes. Because the 11° disk is small enough that both leading and trailing edges fit comfortably within the excitatory region, there is a single peak as the disk passes across the receptive field.

Conversely for the 17° disk, activation is a little weaker at the moment indicated with the arrows in Figure 7E. As we see from Figure 7B, in the left eye the leading edge has just passed out of the central excitatory region into the outer excitatory region where excitation is weaker, while the trailing edge is still in the inhibitory surround. So, the corresponding value of *v_L_*, pointed to by the arrow in Figure 7E, is lower than it was a moment ago. It will shortly rise again when the trailing edge enters the excitatory region. For the 17° disk, there are thus two distinct peaks of activity in *v_L,R_*, which occur when first the leading and then the trailing edge of the disk cross the central excitatory region.

For the largest disk shown, in Figure 7CFI, there are also two peaks, but this time the central dip between them is much lower - in fact slightly negative - because there is a time when both edges are almost entirely within the inhibitory surround (cf right eye in Figure 7C). The peaks on either side of the dip are also lower. This is because the larger disks don’t fit entirely within the excitatory region. When the leading edge is crossing the central excitatory region, the trailing edge is still in the inhibitory surround. Because the weight of the central region is stronger, there is still net excitation, but less than for the smaller disk. In fact, this disk is too large for even one edge to fit within the excitatory region: the top and bottom are always within the inhibitory surround. As expected given its centre/surround structure, the monocular receptive fields are size-tuned, responding maximally to disks around 11 deg in diameter. This is why the peak value of *v_L,R_* is larger in Figure 7D than in Figure 7F.

#### 3.2.3 Binocular response

We now consider the binocular response. The bottom row of panels shows the output of the disparity sensor, *R* = [*v_L_*(*t*) + *v_R_*(*t*) + *b*]^*γ*^ (Equation 2), with the fitted values b = −0.054 and *γ* = 5.05. Instead of plotting this directly as a function of time, we have shown it as a function of *x_c_*(*t*) = 0.5(*x_tgtL_*(*t*) + *x_tgtR_*(*t*)), the visual direction of the target. Again, because the target is moving from left to right at constant speed, these two representations are equivalent. Values of *x_tgtL_*(*t*) and *x_tgt_*, (*t*) for given *t* always differ by the disparity of the target, here 12°. The arrows from Figure 7D to G show two examples. The solid arrows show the value of R when the target crosses the midline, *x_c_* = 0°, while the dotted arrows are for the earlier time when *x_c_* = −6°.

The expected number of strikes for a given trajectory is found by integrating the response along the corresponding row of the panel. Thus, the strike rate depends not only on the peak value, but also on the number of peaks. For the parameter optimisation and for the results shown in Figure 6, the target passed directly over the receptive field center, corresponding to the trajectory with *y_c_* = 0. (As Figure 9 makes clear, if the target does not pass directly over the center, the response is reduced but the dependence on size and disparity remains similar, so the restriction to targets passing directly over the receptive field does not lose behaviour of interest.)

The high value of the output exponent *γ* makes the sensor extremely sensitive to the value of the combined receptive field outputs, *v_L_* + *v_R_*. This enhances the size tuning and also ensures that the sensor is both very sensitive to disparity and unresponsive to monocular stimuli. At the time of the snapshot in Figure 7A, the inputs to the sensor from each eye, *v_L_* and *v_R_*, are each close to their peak value, roughly 0.35. The sensor response is therefore (0.35 + 0.35 – 0.054)^5.05^ = 0.11.

However, if the inputs to the sensor halve, the sensor output falls by a factor of 50 ((0.35 – 0.054)^5.05^ = 0.002). The fairly small reductions in receptive field output as the disk size increases are thus greatly exaggerated in the disparity sensor (compare Figure 7D,E,F with Figure 7G,H,I). The same effect means that there is virtually no response to monocular stimuli, where the peak input is necessarily halved.

#### 3.2.4 Disparity tuning

Figure 8 shows the effect of altering the binocular disparity. It is the same as Figure 7, except for a disk on the screen plane, at a distance of 10cm, instead of at a simulated distance of 2.1cm (and with a different color axis for the binocular response row). The target screen disparity is now zero: that is, the left and right half-images always appear at the same locations on the screen, making them offset from the sensor receptive fields. As is clear from comparing Figure 8 with Figure 7, the monocular time courses are unchanged, but they are shifted relative to one another. The peak response to the 11° disk is thus greatly reduced: when the right-eye input is maximal, the left-eye input is minimal. The maximum input to the binocular sensor is therefore reduced, and the output exponent γ amplifies this further, so the sensor response is very weak (dotted arrows linking Figure 8DE to G). The strongest response is elicited when both inputs are medium (solid arrows) but this response is far weaker than when the target disparity matched the sensor disparity (compare Figure 8G with Figure 7G). Since strike probability reflects the activity of this sensor integrated over time, this weaker response explains the much lower strike rates shown in Figure 6 (70% strike probability for an 11°-disk at 2cm, compared with < 10% at 10cm).

**Fig 8.**
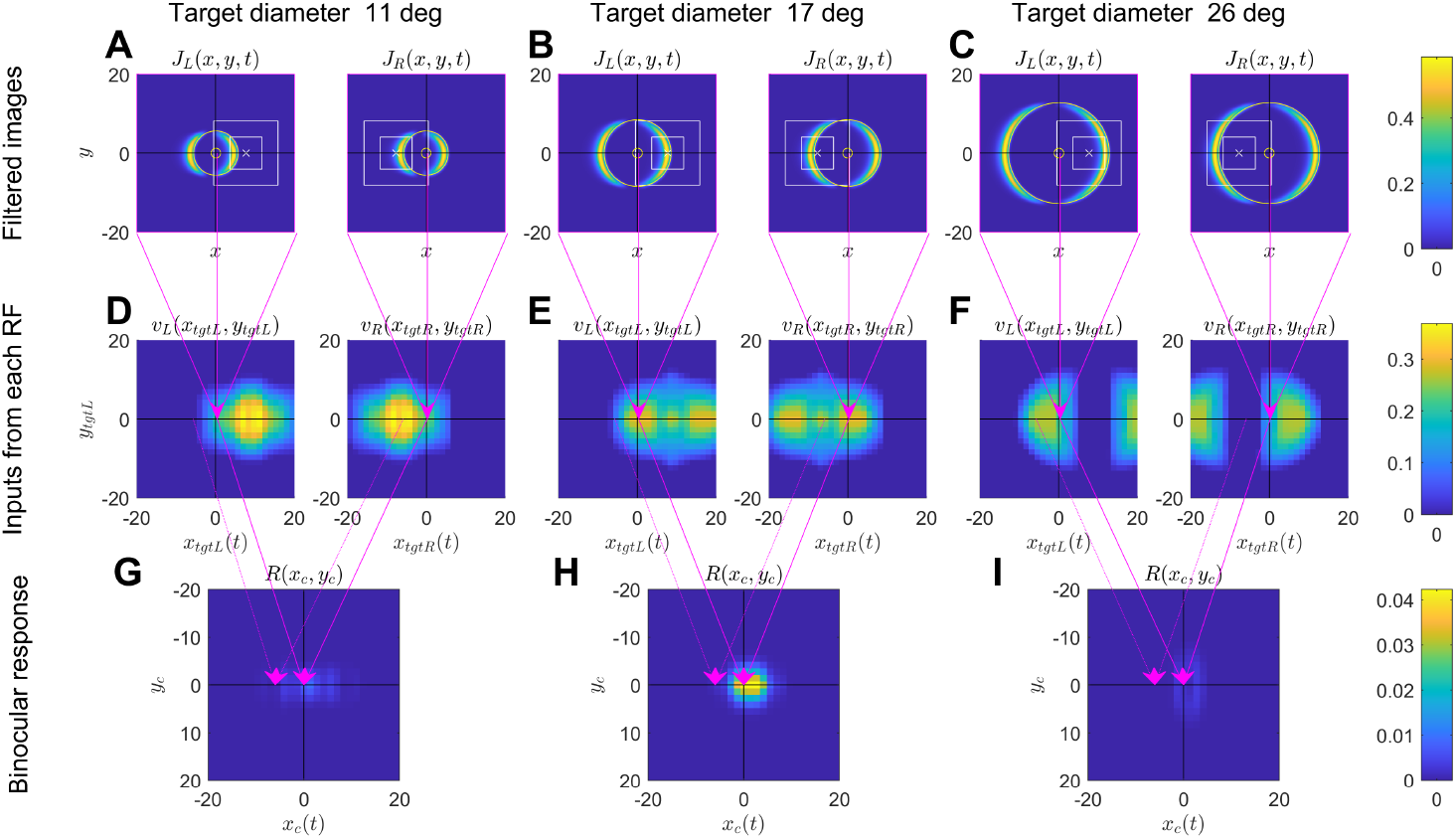
As for Figure 7, but for a target at a simulated distance of 10cm from the mantis (zero screen disparity).

#### 3.2.5 Disparity-dependent size tuning

As discussed above, Figure 6 also shows the interesting property that the preferred target size varies with disparity. Targets 2cm from the mantis produce the most strikes if they are around 10° in diameter. However, targets 10cm from the mantis produce the most strikes if they are around 17° in diameter. Figure 8 explains why this occurs.

For the 17° disk, the separation of leading and trailing edges is close to the difference between the disparity of the sensor receptive fields and that of the target (*α_pref_* = 15.4°). This means that the leading edge in the left eye crosses the central excitatory region of the left-eye receptive field at the same time as the trailing edge in the right eye crosses the central excitatory region of the right-eye receptive field (Figure 8B). Accordingly, the binocular sensor receives strong input from both eyes simultaneously (solid arrows from Figure 8E to Figure 8H). This explains why there is a relatively strong response in Figure 8H at *x_c_* = 0. It is due to a “false match” between the leading edge in the left eye and the trailing edge in the right eye. This response is still weak compared to the response to the optimal size and disparity (note the different colorscale in Figure 8GHI vs Figure 7GHI) but it does explain the shift in size tuning with distance that we see both in the empirical data and in the model (Figure 6). As the target passes in front of the mantis, there is only one false match (leading-trailing), whereas when the target is at 2.1cm there are two true matches (leading-leading and trailing-trailing; cf double peaks in Figure 7H). Thus the true matches always produce the highest activity averaged over time, and so at all sizes, the highest strike rate is obtained when the target is at 2.1cm (the disparity tuning is not size-dependent, Figure 6). But the false match possible for large targets explains why the size-tuning is disparity-dependent.

#### 3.2.6 Effect of direction of motion

False matches between leading and trailing edges also account for the weak but non-zero strike rates predicted for uncrossed stimuli. Because the receptive fields have only horizontal disparity, not vertical disparity, false matches between leading and trailing edges can occur only for horizontal target motion. This explains why the model predicts disparity-dependent size tuning for crossed targets, and strikes to uncrossed stimuli, only for targets moving horizontally (Figure 6A vs B).

Figure 9 shows the sensor response for different sizes, disparities and directions of motion, in the same way as above for Figure 7GHI. The color of each pixel represents the instantaneous response as the disk passes the location (*x_c_, y_c_*). For the disks moving horizontally (Figure 9A-L), *x_c_*(*t*) increases linearly with time whereas *y_c_* is constant; thus each pixel row in a subpanel shows the time-course of the response for a disk at the given y-position. For the disks moving vertically (Figure 9M-X), *y_c_* (*t*) increases linearly with time whereas *x_c_* is constant, so here each pixel column shows the time-course for a disk at the given x-location. Figure 9EI corresponds to the horizontally-moving disks at 2.5cm which were shown in Figure 7GH, with diameters 11 and 17° respectively.

**Fig 9.**
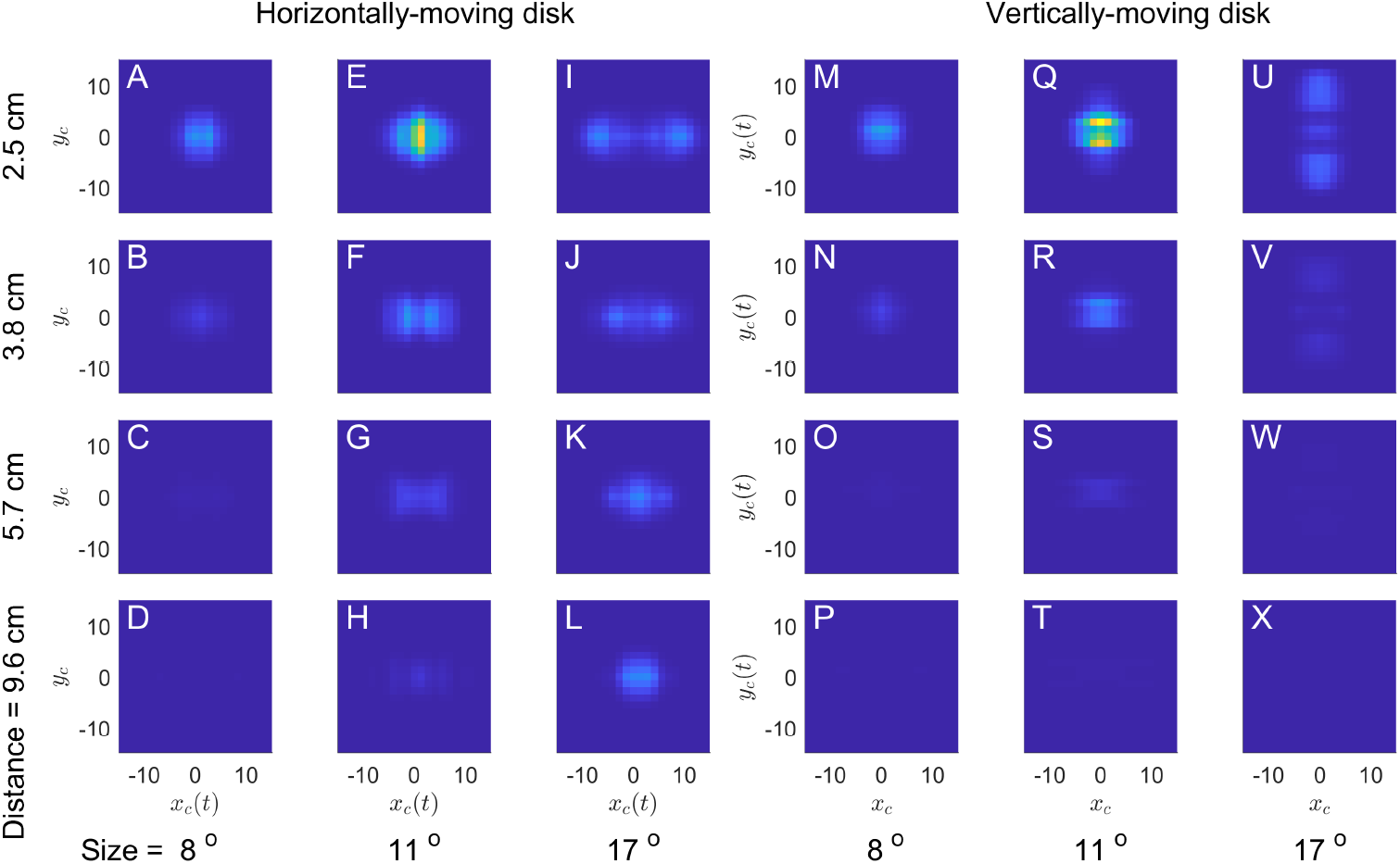
Model sensor output R(x_c_, y_c_) for horizontally or vertically travelling targets as they move across the centre of the field of view. Different rows are for different target distances, as indicated down the left-hand side of the figure, while different target sizes are in columns, as indicated along the bottom. The pseudocolour shows the instantaneous response of the disparity sensor as the centre of the moving target first reaches the location indicated by the (x, y) position. (To simulate the 60Hz refresh rate of the monitor, the target changes position every 16ms, and so stays in each location for five of the simulation’s 3.3ms timesteps.) The color axis is the same in all 24 panels

For a disk at 2.1cm, corresponding to the sensor disparity *d* (Table 1), the response would be identical for both directions of motion. This is because in this case, the target passes exactly through the middle of the receptive field in each eye, at the same time in both eyes. Because both the target and receptive fields are unchanged by rotation through 90 degrees, the response is also unchanged. The top row of Figure 9 shows responses for disks at 2.5cm, close to the sensor disparity, and indeed the response as a function of time is very similar for the two directions of motion (though of course rotated as a function of (*x_c_, y_c_*) to reflect the direction of motion).

For other distances, this is not so. For stimuli at greater distances, the stimulus horizontal disparity is smaller and thus more mismatched with the sensor horizontal disparity. This reduces the response for both directions of motion, but the underlying reason is different depending on the target direction of motion, as we now explain.

For vertical motion, assuming that the target has no vertical disparity, the leading and trailing edges enter the excitatory region at the same time, but if the target disparity does not match the sensor disparity, they cannot both pass over its center. If the target has a suitable horizontal offset *x_c_*, it can pass over the optimal region in one eye, but in the other eye much of the target necessarily passes over the inhibitory region, causing a weak response. The best compromise is for *x_c_* close to 0, when the target is only slightly misaligned in both eyes. Since the sensor receptive fields are symmetrically located about the midline, the sensor is tuned to stimuli on the midline, and so it responds most strongly to stimuli on the midline even when these do not have the preferred disparity. This is why the peaks are centred on *x_c_* = 0 in Figure 9M-X, the panels relating to vertical motion.

For a target moving horizontally (and with no vertical disparity), the target can pass over the central excitatory region in both eyes. However, if the target disparity does not match that of the sensor, the leading and trailing edges are offset relative to the receptive fields: compare Figure 7ABC with Figure 8ABC. This usually reduces the response.

However, for some combinations of target size and disparity, the leading edge can pass through the excitatory region in one eye at the same time as the trailing edge is passing through the excitatory region in the other eye, as in Figure 8B. The two mismatched edges thus combine to give a response in the disparity sensor comparable to the response in the top rows of Figure 9, where both leading edges crossed at the same time. This effect occurs when the difference between the target and sensor disparity is similar to the target diameter. For example, for a target at a distance of 10cm, the target disparity is 4 deg, 11.4 deg less than the sensor disparity of 15.4 deg. For a target of diameter 17deg, this is close enough for the leading and trailing edges to overlap substantially. This explains why a 17-deg diameter is the only size shown in Figure 9 where a target at 10cm still elicits a weak response (panel L).

### 3.3 Predicted striking behaviour as a function of vertical disparity

Rossel and colleagues [31] examined praying mantis striking behavior in response to stimuli with vertical (non-epipolar) disparity, introduced by means of prisms. They found that the sensitivity to horizontal disparity was unchanged, but that the strike rate declined with vertical disparity, reaching zero when the vertical disparity exceeded 15°. Strike rate depended on size, as already noted, but the dependence on vertical disparity was independent of size. The authors concluded that “the limit for vertical disparities is fixed, regardless of the size and microstructure of the target”.

We examined the response of our disparity sensor to stimuli with vertical disparity. Figure 10 shows the mean predicted strike rate, *M_model_* (Equation 3), for targets moving horizontal and vertically, as a function of vertical disparity Δy (shown on the horizontal axis). The vertical axis shows the offset of the target trajectory from the receptive field center, measured perpendicularly to the direction of motion. The different panels show results for targets of different sizes and distances.

**Fig 10.**
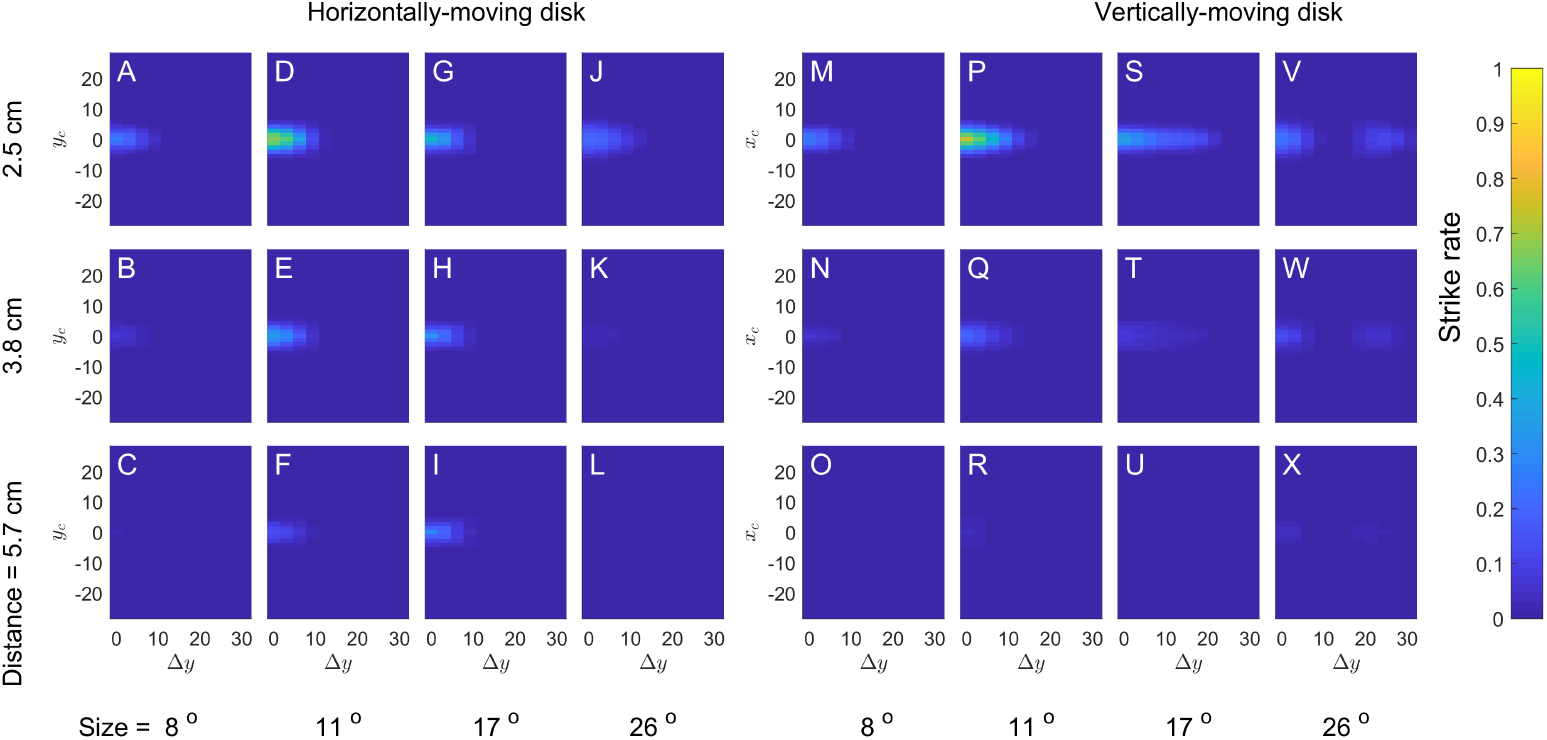
Predicted strike rate M_model_ (Equation 3) for disks moving horizontally (A-L) or vertically (M-X) in front of the model mantis, as a function of vertical disparity Δy and offset perpendicular to the direction of motion (y_c_ for horizontally-moving targets and x_c_ for vertically-moving). For the horizontally-moving targets, the vertical location of the target in the two eyes is constant at y_L_ = y_c_ + Δy/2 and y_R_ = y_c_ – Δy/2. For vertically-moving targets, the mean y-location is of course a function of time, but the difference y_L_ – y_R_ is constant at Δy. Different rows are for different disparities, thus simulating disks at different distances from the animal when Δy = 0. Different columns are for different disk diameters.

For all sizes and distances, the highest strike rate occurs for stimuli with zero vertical disparity and zero offset (i.e. passing directly over the receptive fields). In all cases, the strike rate falls to zero for vertical disparities of around 15°, corresponding to the extent of the excitatory region. For vertical disparities greater than this, it is impossible for the target to pass over the excitatory regions in both eyes simultaneously. However for large vertically moving disks that are close to the sensor’s preferred disparity, a low but non-zero strike probability is observed for very large vertical disparities. This is again the result of a “false match”, when the target’s trailing edge in one eye passes over the excitatory region at the same time as the leading edge in the other one.

Figure 11 plots the strike rates predicted by the model for zero offset as a function of vertical disparity (solid lines), and compares them with the empirical results of [31] (dashed lines).

**Fig 11.**
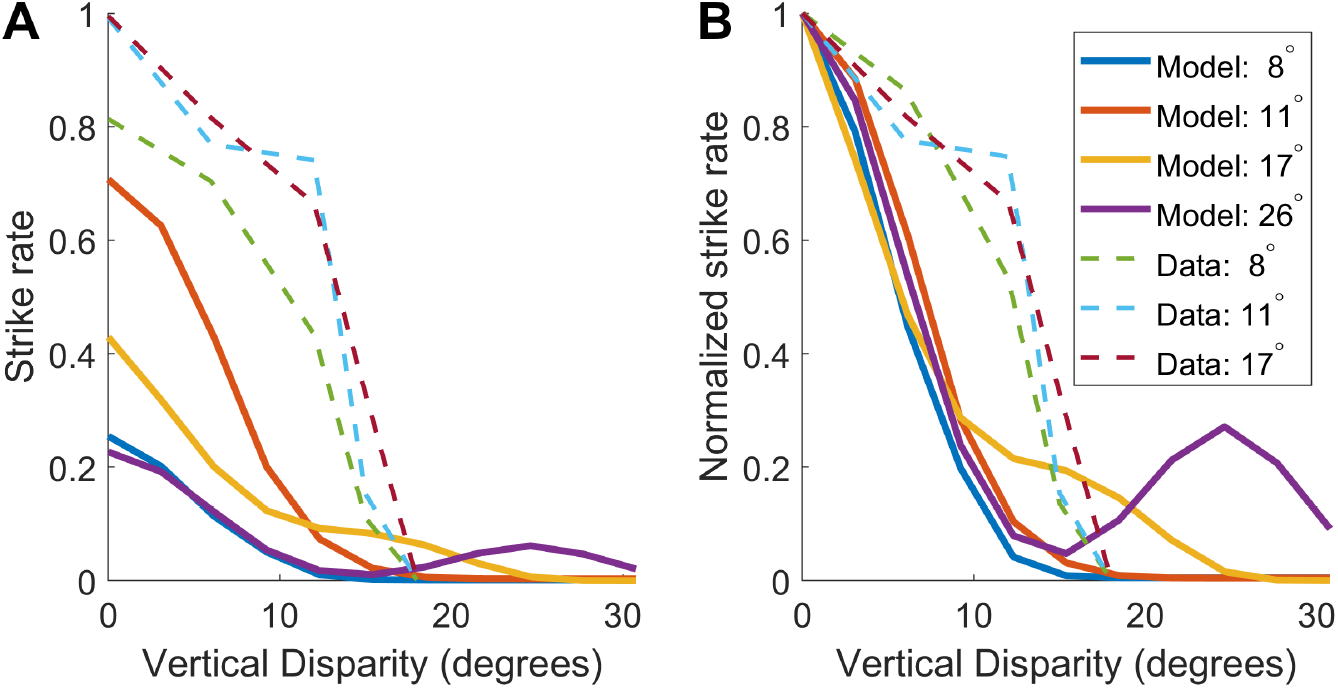
Sensitivity to vertical disparity and size, for our model (solid lines) and empirical data digitized from [31]. A: Solid lines show strike rate predicted by our model, averaged across horizontally and vertically travelling targets passing directly over the sensor. Dashed lines show strike rates observed by [31] for stationary “jiggling” targets. Colors indicate target size as shown in the legend. B: as A but the strike rate is normalized to be 1 for stimuli with zero vertical disparity

## 4 Discussion

In this paper we set out to find out how well the praying mantis’ strike response to stimuli of different sizes and distances could be captured using a single disparity sensor. The binocular computation carried out by this model sensor is closely based on the stereo energy model. However, the early processing is different, incorporating lowpass spatial filtering, highpass temporal filtering and squaring. The model has 8 free parameters (Table 1): 5 which describe the size and weights of the monocular receptive field functions (*s*_*e*1_, *s*_*e*2_, *w*_*e*1_, *w*_*e*2_, *w_i_*), two describing binocular combination (*b* and *γ*) and one describing the preferred distance (*α_pref_*). We fitted these parameters to our empirical data [8] on strike rates for moving targets of a range of sizes and simulated distances. We also examined the predictions of this model to stimuli with vertical disparity.

We were able to find parameters that enabled this sensor to capture the key features of interest (Figure 6): (1) strong tuning to disparity (strike rate reduced by a factor of 4 for a 5° change in disparity); (2) weak tuning to size (strike rate reduced by a factor of 4 for a 20° size in diameter. This confirms that our data can be modelled with a single sensor, tuned to both disparity and to size.

Our model is also qualitatively consistent with Rossel’s data on vertical disparity (Figure 11). Our data were not fitted to these results, which were from a different species with a larger size preference, so in our model the strike rate falls to zero at a smaller size than in Rossel’s data. However, the model shows the same phenomenon reported by Rossel: that the fall-off in strike rate with vertical disparity was independent of the stimulus size (Figure 11B), so that it is not the case that vertical disparity is tolerated so long as the stimuli in the two eyes overlap by a certain extent. Our model makes it clear that the limit on vertical disparity is set by the receptive field size, and thus by the animal’s preferred size rather than the size of the particular stimulus. The model thus proposes a neurophysiological explanation for Rossel’s finding of a fixed limit for vertical disparities.

Our model also produced three other surprising results, all due to false matches between the leading and trailing edges of the stimulus. First, for horizontally-moving stimuli, the model prefers larger angular sizes at further distances. This effect is seen in our data [8], though since we used spiralling stimuli we cannot say whether it occurs only for horizontal trajectories. It was a puzzling result since it is the opposite of what would be required for physical size constancy We had not considered the possibility that false matches between leading and trailing edges could account for it. Second, also for large horizontally-moving stimuli, the model predicts a low but non-zero strike rate for uncrossed disparities, which do not correspond to a single point in space. We saw a similar effect in our data but argued that it was not driven by disparity, so subtracted it before fitting. We now wonder if these strikes could have been driven by false matches in the mantis as they were in our model. Finally, the model predicts that vertically-moving stimuli, if much larger than the preferred size, can elicit strikes even when they have very large vertical disparity Figure 11. Since the only experiments with vertical disparity used stationary stimuli [31], we do not know whether such an effect would be observed empirically.

For uniform light stimuli like those used here, the leading edge represents a contrast increment while the trailing edge is a decrement. A false match between these two is only possible because our model includes a rectification step which abolishes the distinction between increments and decrements. Neurally, this would correspond to combining inputs without regard to whether they indicate increments and decrements, e.g. excitatory input from both L1 and L2 pathways [27]. An alternative would have been to postulate input only from the L2 pathway, which responds to moving dark edges. Mathematically, this would correspond to halfwave instead of full rectification, and would have the advantage of building in mantids’ preference for dark prey items [22], currently ignored in our model. However, this would have meant that the sensor would “see” only the leading edge of the target, and so would be insensitive to size parallel to the direction of motion, which is not what is observed. Our decision to make the sensor respond to both leading and trailing edges enabled us to achieve realistic size-tuning, but enables these false matches. It therefore becomes important to test these predictions of the model behaviorally, in order to understand whether mantis stereopsis can really be misled by false matches between the leading and trailing edges of a moving target, or whether mantids have a more sophisticated form of stereo correspondence which prevents this.

The above discussion raises the question of what aspects of mantis striking behaviour should be attributed to the properties of their disparity sensors, and what should be considered separate. The model discussed in this paper is sensitive to size and distance, but - as just noted - not to contrast polarity, nor directly to other aspects of visual stimuli that influence strike rates, such as looming, speed of figure motion etc. We have argued previously [12] that mantis behaviour is most economically accounted for by postulating neural mechanisms for detecting prey which are distinct from those detecting binocular disparity: effectively, a “prey sensor” in addition to the disparity sensor modelled here. We proposed that striking requires the disparity sensor to be activated at the same time as, or within a limited time window after, the prey sensor, and that the strike rate reflects the activity in both systems (e.g. via a multiplicative interaction). We argued that this can explain why looming (an increase in angular size over time) increased strike rates independent of disparity [32]: looming activates the prey sensor but does not enhance the activity of the disparity sensor. It also explains why mantids can discriminate stereoscopic depth in stimuli with no figure motion, although they will not strike at these unless they first see stimuli with figure motion [12]: the figure motion activates the prey sensor, and if the disparity sensor is subsequently activated as well, strikes can occur. In principle, we could have attributed the size tuning also to properties of the prey sensor, but instead we chose to incorporate it within the disparity sensor directly. Two lines of evidence, one behavioural and one neurophysiological, motivated this decision. First, neurons sensitive to disparity have centre/surround receptive fields [13, 14], implying that they are tuned to size as well as to disparity. Second, the dependence of size tuning on disparity [8] suggests joint rather than separable mechanisms.

Our model disparity sensor is not tuned directly to speed or figure motion, but its temporal filtering makes it indirectly sensitive to target speed. This is because the speed affects the width of the leading and trailing edges across the retina. The squared output of the high-pass filter is above 14% of maximum for a time *τ* after the edge passes, so the width is *V_τ_*. Slower stimuli thus offer less total excitation as they pass over the receptive field. Faster stimuli give wider edges, but once the stimulus gets fast enough that the leading and trailing edges merge, there is less total excitation. All in all, these effects lead to complicated interactions between speed, size and disparity tuning which are beyond the scope of this paper. We do not, however, believe that the current model is accurate in this regard. More accurate models of the spatiotemporal inputs to the disparity sensor will be required to ensure that model predictions remain accurate for stimuli moving at different speeds, and for stimuli more complicated than black disks (e.g. targets defined by luminance flicker or theta motion [11, 12]). This will require more detailed behavioral data, and would benefit greatly from further neurophysiological experiments elucidating the properties of the relevant neurons.

Finally, this model includes only a single disparity sensor. As noted in the Introduction, it is helpful to begin with the simplest possible model in order to understand what this is capable of, before elaborating it. In fact, neuroanatomical and physiological data suggests multiple disparity-tuned neurons, even within a given cell class [13], as well as multiple classes of disparity-tuned neuron whose role is unclear. Some may mediate other behavior, such as head saccades, which are also sensitive to stereoscopic information; others may implement top-down feedback, for example guiding visual attention in space. One simple extension of the present model would be to include a few different copies of this sensor, with receptive fields at different locations in the visual field. This would account for mantids’ ability to strike at targets over a range of the visual field around 20° in diameter. However, expanding the model to account for the full range of disparity tuning found in mantis brain must await more neurophysiological data on their response properties.

